# Fast and global detection of periodic sequence repeats in large genomic resources

**DOI:** 10.1101/309039

**Authors:** Hideto Mori, Daniel Evans-Yamamoto, Soh Ishiguro, Masaru Tomita, Nozomu Yachie

## Abstract

Periodically repeating DNA and protein elements are involved in various important biological events including genomic evolution, gene regulation, protein complex formation, and immunity. Notably, the currently used genome editing tools such as ZFNs, TALENs, and CRISPRs are also all associated with periodically repeating biomolecules of natural organisms. Despite the biological importance of periodically repeating sequences and the expectation that new genome editing modules could be discovered from such periodical repeats, no software that globally detects such structured elements in large genomic resources in a high-throughput and unsupervised manner has been developed. Here, we developed new software, SPADE (Search for Patterned DNA Elements), that exhaustively explores periodic DNA and protein repeats from large-scale genomic datasets based on *k*-mer periodicity evaluation. SPADE precisely captured reported genome-editing-associated sequences and other protein families involving repeating domains with significantly better performance than the other software designed for limited sets of repetitive biomolecular sequences.

---

Significant roles of repetitive DNA and protein sequences have been widely reported in both eukaryotes and prokaryotes. Transposable DNA elements are thought to be among the most important evolutionary driving factors that have been expanding within and between species’ genomes via their copy-and-paste or cut-and-paste mechanisms^1,2^. These repetitive elements induce large-scale genomic rearrangements and transcriptional regulation. Nonmobile short tandem repeat DNA sequences are also key elements inducing structural genome evolution in prokaryotic species. Tandem DNA repeats induce mispairing and slippage between repetitive units during DNA replication and drive genomic contraction and expansion^3^. Intramolecular crossover DNA recombination is also promoted between tandem repeat regions of the genome^4^. Some of these events are known to be reversible and lead to genomic phase variation, allowing cells and species to rapidly adapt to changing environments without having to undergo irreversible mutations^5^.

Repetitive protein domains often serve as structural binding modules that stably interact with biopolymers. They are also widely involved in protein folding and interactions in various biological processes. Tetratricopeptide repeats (TPRs)^6^ and ankyrin (ANK)^7^ repeats are large protein repeat families that are conserved from prokaryotes to eukaryotes. The repeat units of TPRs and ANK repeats are 34 and 33 amino acids (aa) long, respectively, and both are composed of a helix–turn–helix structure. These repetitive domains have been reported to mediate interactions with other proteins and RNAs and play important roles in cell cycle control, transcriptional regulation, translational inhibition, and protein translocation^6,7^. WD40 repeat is another large protein repeat family found in both prokaryotic and eukaryotic species, but its functions are particularly well known in eukaryotes^8^. A WD40 repeat is composed of seven-bladed β-propellers, where each propeller is around 40 aa long, involving four antiparallel β-sheets, and serves as a scaffold for protein interaction. Accordingly, WD40 proteins coordinate multi-protein complex formation and underlie diverse biological functions such as signal transduction, transcriptional regulation, cell cycle control, chemotaxis, autophagy, and apoptosis^8,9^.

The structural code for proteins in general remains largely unclear and there have been major challenges in engineering these repeat protein modules to develop synthetic binding reagents for biomedical and nanotechnology applications^10,11,12^. Most protein repeat sequences are “imperfect” or “degenerated,” where each repetitive unit contains variable amino acid residues and the degrees of repeat imperfectness vary widely. Some of these variable residues determine binding to specific biomolecules and deciphering this code for DNA binding has been extremely beneficial in the development of genome editing technologies^13,14^. Several transcriptional regulators involve tandem protein repeats with specific periodicities to wrap around the double-stranded DNA helix. Cys2His2 zinc fingers (C2H2 ZNFs) are the most common DNA-binding motif found in eukaryotic transcription factors^15^. C2H2 ZNFs represent periodic protein repeats that make tandem contact to targeting DNA sequence. The repeat unit size ranges from 28 to 30 aa and the variable amino acid residue pattern in each unit defines its binding to a specific DNA triplet^16^. Similarly, transcription activator-like effectors (TALEs) of the type III secretion system encoded in the plant pathogenic bacteria of the *Xanthomonas* genus also have repeating domains^17^. They are virulence proteins that bind to the host plant genomic DNA and hijack its gene expression system. The periodicity of the repeat unit ranges from 33 to 35 aa, where the combination of two variable amino acid residues at the 12^th^ and 13^th^ positions of the repeat sequence has a one-to-one relationship with a specific mononucleotide. By fusing DNA cleavage domains such as FokI endonuclease to C2H2 ZNFs and TALEs, the genome editing tools zinc finger nucleases (ZFNs) and TALE nucleases (TALENs), respectively, have been developed, both of which enable highly specific targeted DNA cleavage. Other effector proteins have also been fused to C2H2 ZNFs and TALEs to regulate gene expression and chromosomal structures in various organisms^18,19^.

The CRISPR–Cas systems have become the most widely used genome editing technologies in recent years^20^. As indicated by their name, clustered regularly interspaced short palindromic repeats (CRISPRs) are widely encoded in prokaryotic genomes^21^. The unique characteristics of these CRISPRs and CRISPR-associated (Cas) proteins in bacterial and archaeal immunity have been rapidly identified recently^22^. In the immunization process, a fragment of defined length from invading phage or plasmid DNA is incorporated into the 5’ end of a CRISPR locus with a constant motif sequence. Accordingly, the periodic interspaced repeats of CRISPRs have been derived by continuous cycles of this immunization process. In the immunity process, an RNA originating from the immunized DNA is transcribed and processed and guides Cas protein(s) to its complementary sequence of exogenous DNA for cleavage and degradation. Harnessing different Cas proteins and RNAs involved in the immunization/immunity processes of different CRISPR-type families, various genome editing technologies have been established^20,23,24^. Cas9 with double-stranded DNA cleavage activity from the type II CRISPR system has been widely used for targeted gene disruption and targeted fragment knock-in in various organisms including mammals. Similar to ZFNs and TALENs, nuclease-deficient Cas9 (dCas9) or mutant Cas9 nickase (nCas9) fused to effector proteins such as transcription factors, deaminases, and fluorescent proteins have been used for various applications such as gene silencing^25^, activation^26^, single-base editing^27^, and chromosomal labeling^28^.

Periodically repeating DNA and protein sequences have diverse and important roles in biology. A simple and optimistic hypothesis has been proposed that new genome editing modules can be discovered from other periodic repeats in large-scale genomic resources. However, there is no universal software that captures various types of periodic repeats from large-scale genomic datasets in an unsupervised manner (Table 1). For example, RepeatMasker is one of the most commonly used tools to detect interspersed DNA repeats and low-complexity DNA sequences^29^. However, this software screens only DNA sequences against a database of reported elements and does not evaluate repeat periodicity. Previous software programs developed for *de novo* searches of repetitive biomolecular sequences also have certain limitations. Tandem Repeat Finder is one of the first types of software to screen tandem and low-complexity DNA repeats without prior knowledge^30^, but is incapable of capturing highly degenerated or interspaced DNA sequences or protein repeats. RECON^31^ and RepeatScout^32^ also screen only DNA sequences, focus only on interspersed repeats regardless of periodicity, and exclude tandem or low-complexity repeats. PRAP captures both tandem and interspersed repeats, but screens only DNA sequences^33^. Although the recently developed software XSTREAM^34^ and T-REKS^35^ search for both tandem and highly degenerate repeats from DNA and protein sequences, both are ineffective at capturing interspersed or interspaced repeats including CRISPRs. With the recent interest in genome editing, several software packages such as CRISPRFinder^36^, CRISPRdigger^37^, and AnnoTALE^38^ have been developed to capture genome-editing-associated sequences. However, such specialized software does not have the potential to discover novel genome editing modules.

**Table 1.**
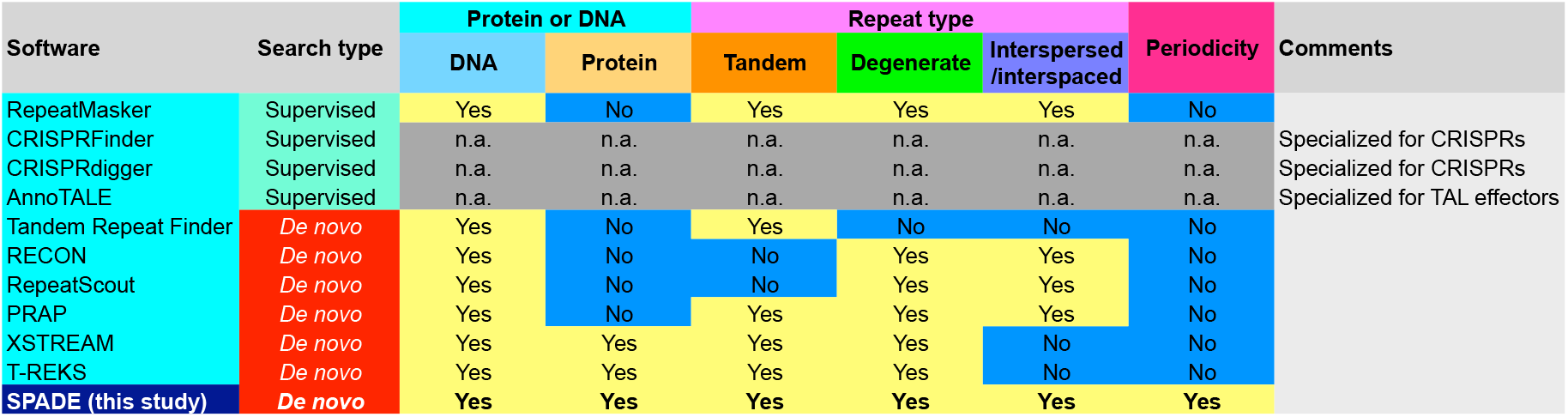
Comparison of different software for capturing repetitive biomolecular sequences

The previously developed software focuses on limited types of repeat sequences for specific biological targets, but it seems that any software that combines the abilities of the previous software packages for any type of repetitive sequence would give an ambiguous and large set of sequences, which would require substantial effort for further curation and validation. However, none of the abovementioned software screens repetitive sequences based on sequence periodicity that commonly appears in many significant biological processes. This could be a strong constraint in screening to obtain a set of biomolecular sequences with high potential for expanding our biological knowledge and developing new biotechnologies. Accordingly, we have been motivated to develop simple and fast software called SPADE (Search for Patterned DNA Elements) that globally captures such periodically repetitive biomolecular sequences in large genomic datasets mainly based on an evaluation of k-mer periodicity.

## Overview of SPADE

We implemented SPADE to efficiently screen periodically repeating sequences as follows (Supplementary Fig. 1). The software first automatically extracts multiple sequence entries from an input file (GenBank or FASTA format) and identifies the sequence type (DNA or protein) for each entry. Each entry sequence is scanned by a sliding window to count k-mers and highly repetitive regions are extracted. The sequence periodicity of each highly repetitive region is then evaluated based on a position-period matrix that cumulatively plots the distance between the same neighboring k-mers and their sequence positions (see Methods). The periodic sequence region is defined, and the periodic sequence units are queried for a multiple alignment to identify repetitive motifs. A representative motif sequence is then aligned back to the entry sequence to annotate the periodically repeating units. Finally, the annotations for the detected periodic repeats are added to the input information and output in the GenBank format with options to visualize *k*-mer density, position-period matrix, repetitive unit loci with neighboring genes, and motif sequence logo for each periodic repeat.

## Periodic repeats in a CRISPR-encoding genome

Using SPADE, we exhaustively searched for periodic DNA and protein sequences in the 7,006 complete prokaryotic genomes that were available in the NCBI RefSeq database. The default parameter set was used for the entire analysis of this study. In the *Streptococcus thermophilus* LMD-9 genome, 7 periodic DNA repeats and 27 periodic protein repeats were detected, including 2 previously annotated CRISPR loci (Fig. 1a and Supplementary Table 1). The repeat periods of the annotated CRISPRs were both 66 bp and their detected repeat motif sequences were identical to the reported motifs (Fig. 1b). Notably, we found a novel interspaced repeat region containing four repeats with a period of 72 bp, in which the repeat motif and interspace sequences were all 36 bp long (Fig. 1c). While type II-A Cas genes were found in the neighboring regions of the reported CRISPRs, a type III-A Cas gene cluster was found in the adjacent region of the novel repeat, suggesting a functional type III-A CRISPR system in this genome.

**Figure 1.**
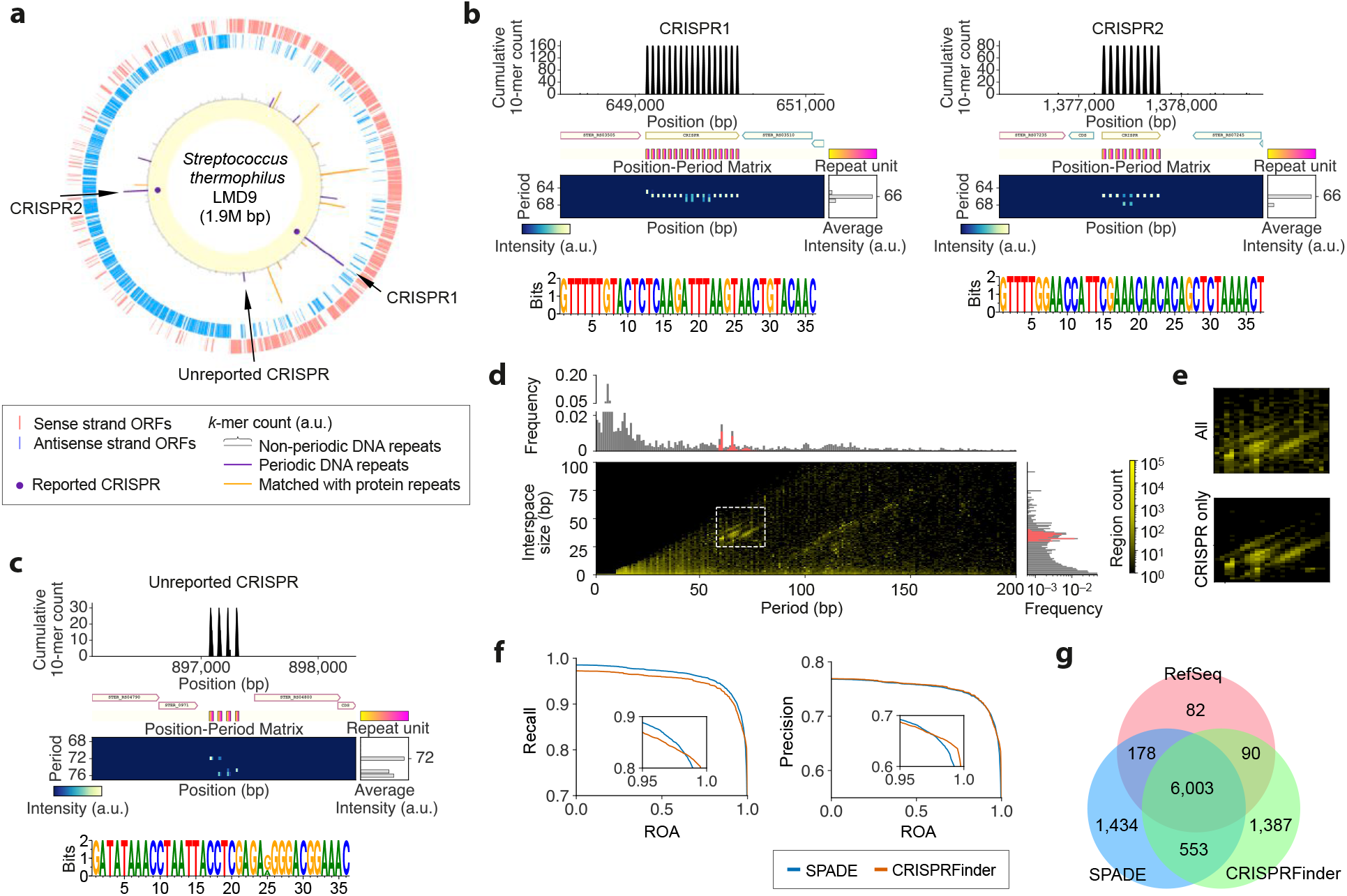
CRISPRs detected by SPADE. (**a**) Circular genome map of the *S. thermophilus* LMD-9 genome. From the outer side, it represents genes encoded on the sense and antisense strands of the genome, cumulative k-mer counts with the annotations of periodic DNA and protein repeats, and the previously reported CRISPR loci. (**b**) Previously reported CRISPRs detected by SPADE. Each periodic repeat region is visualized along with cumulative *k*-mer count, neighboring genes, positions of repeat unit sequences and position-period matrix of the surrounding genomic region, and its motif sequence is represented by sequence logo. Each periodic repeat unit is represented by a gradient box where color indicates relative position in each repeat unit sequence, (**c**) A novel CRISPR found in the *S. thermophilus* LMD-9 genome, (**d**) Period-interspace size distribution of the entire periodic DNA repeats captured in the 7,006 RefSeq prokaryotic genomes. Magenta bars in the probability density distributions represent CRISPRs reported in the RefSeq dataset. Dashed white line box represents DNA repeats further screened as CRISPR candidates, (**e**) Enlarged view of the dashed white line box in (**d**) and distribution of the RefSeq CRISPRs in the same area, (**f**) Precision and recall in predicting RefSeq CRISPRs by SPADE and CRISPRFinder along with region overlap agreement (ROA) thresholds, (**e**) Venn diagram for DNA repeats detected by SPADE and CRISPRFinder with ROA of ≥50% and their agreement with RefSeq CRISPRs.

The other periodic DNA repeats were all short tandem repeats with a period size of 1–7 bp that were commonly found in prokaryotic genomes (Fig. 1d). Among the 27 periodic protein repeats, 24 were short tandem repeats with periodicity of 10 aa or less. The other three included a peptidoglycan-binding protein (three repeats with a 17-aa period) and a subtilisin-like serine protease (three repeats with a 32-aa period), both of which were annotated to involve protein repeats, and a nucleotide exchange factor (four repeats with a 14-aa period), which was annotated to involve two α-helices.

## Performance in detecting CRISPRs

We then measured the performance of SPADE in detecting CRISPRs, the annotation criteria of which are standardized in the NCBI prokaryotic genome annotation pipeline^39^. From the entire 161,465 periodic DNA repeats detected in the 7,006 prokaryotic genomes, we obtained 8,168 genomic regions with a repeat period size and interspace size of 58–81 bp and 25–60 bp, respectively (Supplementary Table 2). These parameters were partly derived from CRISPRFinder, the most commonly used tool for CRISPR annotations in recent genomic resources^36,40,41^, and partly defined empirically based on the reported CRISPRs (see Methods). We confirmed that the distribution of period-interspace size combinations for the defined parameter space had good agreement with that for the 6,354 reported CRISPRs in the RefSeq database (Fig. 1d and e). We then compared the performance of SPADE and CRISPRFinder in capturing CRISPRs. In the same genomic datasets, 8,033 regions were detected by CRISPRFinder (Supplementary Table 2). Precision and recall were decreased along with region overlap agreement (ROA) with reported CRISPR regions for both SPADE and CRISPRFinder, but recalls by SPADE were higher overall than those by CRISPRFinder for ROA of up to 98%, while the levels of precision were similar for the two types of software (Fig. 1f). This indicated that SPADE was slightly better at roughly capturing CRISPRs, but not at the single-base resolution. At 50% ROA, SPADE and CRISPRFinder captured 6,181 and 6,093 RefSeq CRISPR regions, respectively, where 6,003 were captured by both types of software (Fig. 1g). In summary, although SPADE was not specifically designed for CRISPR annotation, its performance for capturing CRISPRs with simple size thresholds was at least on par with the most commonly used CRISPR prediction software.

## Periodic repeats in a TALE-encoding genome

In the *Xanthomonas oryzae* pv. *oryzae (Xoo)* PXO83 genome encoding TALE genes and TALE pseudogenes^38,49^ DNA repeats and 194 protein repeats were detected by SPADE (Fig. 2a and b and Supplementary Table 3). All of the reported TALEs were recaptured with a repeating period of 34 aa and variable residues at the 12^th^ and 13^th^ amino acid residues and two α-helices in each repeat unit, which were all consistent with the reported features of TALE. In the intergenic genomic regions, the previously identified TALE pseudogenes TalAI3 and TalAI4 were also both detected with a period of 102 bp, which was concordant with the period of 34 aa for TALEs (Supplementary Fig. 2). We also detected an annotated large CRISPR locus where a highly constant motif of 31 bp was repeated periodically 86 times each with an interspace sequence of around 34 bp (Fig. 2c).

**Figure 2.**
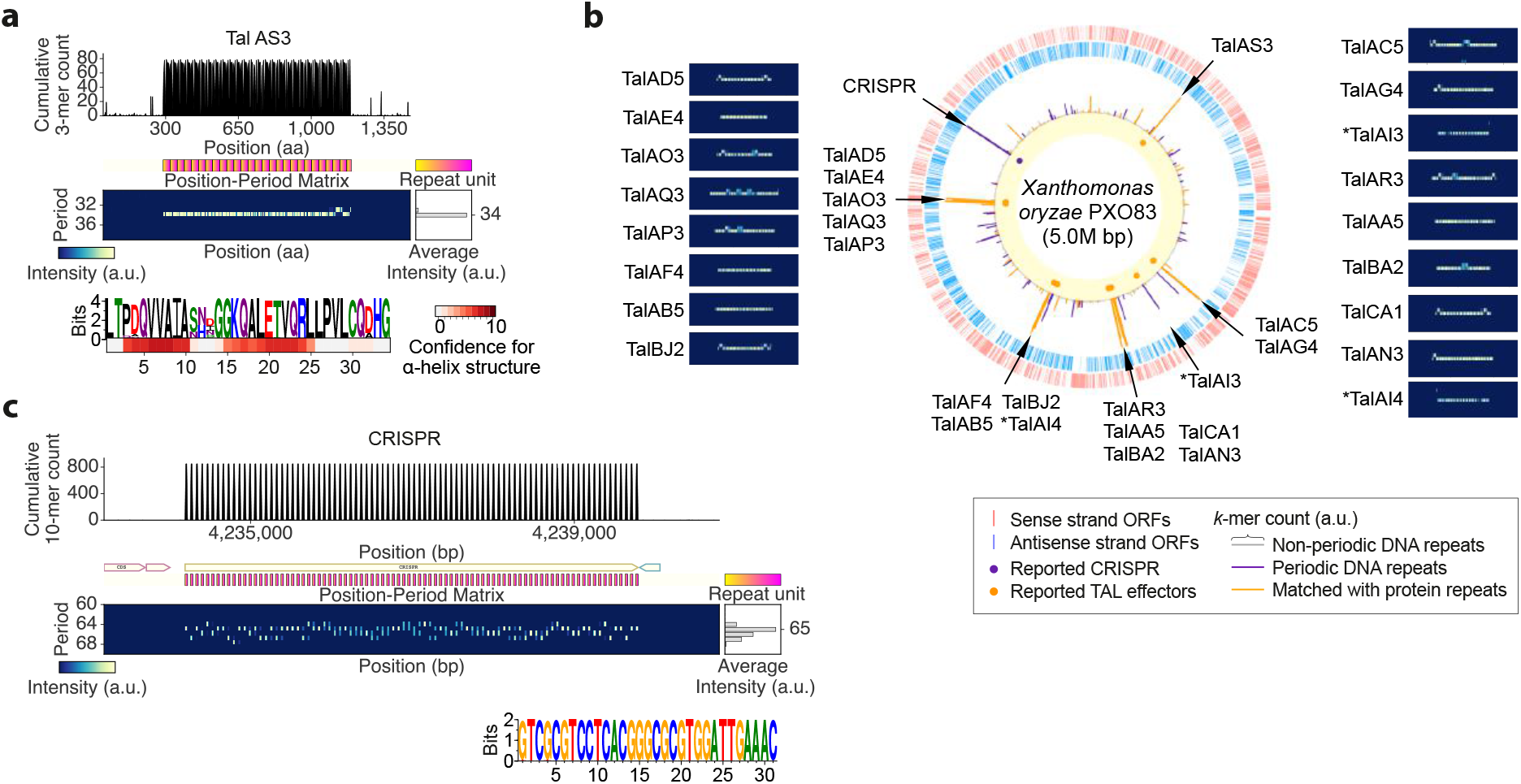
TALEs and a CRISPR detected in the *Xoo* PXO83 genome. (**a**) Example (TalAS3) of TALEs detected by SPADE. For details, see Fig. 1b. The heat map under the repeat motif sequence logo represents confidence scores for α-helical structure at each amino acid residue. (**b**) Circular map representation of the *Xoo* PXO83 genome along with position-period matrices for all of the TALE genes and TALE pseudogenes (denoted by stars) annotated in the RefSeq database. For details, Fig. 1a. (**c**) CRISPR locus of the *Xoo* PXO83 genome detected by SPADE.

Among the other 46 periodic DNA repeats, 40 were short tandem repeats with a period of 10 bp or less, including 25 heptamer repeats that were previously suggested to contribute to phase variation in the *Xanthomonas* genus^42^. Three short tandem DNA repeats were found in intergenic regions, one with a period of 12 bp and two with a period of 14 bp. Another short tandem DNA repeat region was found in the middle of an ABC transporter-encoding gene with a period of 16 bp, which is relatively prime to 3, the protein coding frame size (Fig. 3a), and another longer sequence with a period of 60 bp was also found to encode a hypothetical gene in less than half of its region (Fig. 3b). Furthermore, we found a large periodic DNA region from the genomic position of 3,559,997 to 3,563,142 (3,144 bp long) with an average period of ~787 bp (Supplementary Fig. 3). Following a transposase-encoding gene, this region involved three different hypothetical genes, each of which was in a different repeat unit. Interestingly, all of these three repeats partially overlapping with protein-coding regions were found to be widely conserved in the *Xanthomonas* genus with different numbers of repeats, but the coding gene architecture had markedly diverged evolutionarily (Fig. 3a–c), indicating that phase variations of protein-coding patterns for these regions rapidly occurred after speciation by genomic contraction and expansion via the repetitive sequences.

**Figure 3.**
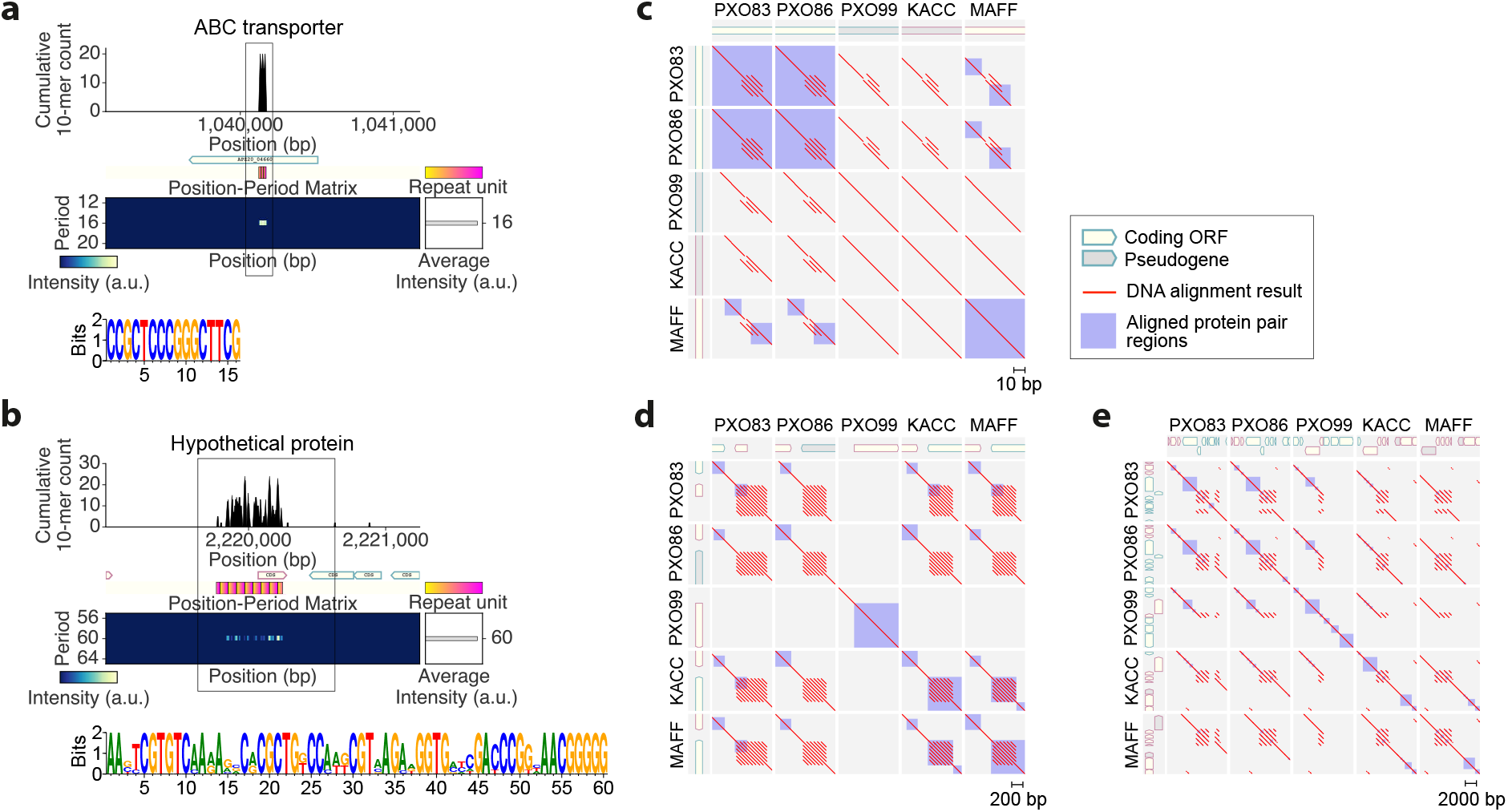
Comparative analysis of the repetitive DNA regions determined to overlap with protein-coding regions in the *Xanthomonas* genus. (**a**) An ABC transporter-coding region involving a DNA repeat of 16-bp period. Line box denotes the genomic regions used for the comparative genomic analysis presented in (**c**). (**b**) A hypothetical protein-coding region overlapped with a DNA repeat of 60-bp period. Line box denotes the genomic regions used for the comparative genomic analysis presented in (**d**). (**c**–**e**) Sequence alignment analysis of the repetitive DNA sequences and their protein-coding patterns amongst the *Xoo* PXO83, PXO86, PXO99, KACC10331, and MAFF311018 genomes. With protein/pseudogene-coding structures under the label of each genome, the within- and between-genome alignment results are represented by red lines for DNA and blue boxes for proteins, respectively. (**c**) For an ABC transporter-coding region in the *Xoo* PXO83 genome. (**d**) For a hypothetical protein-coding region overlapping with a DNA repeat of 60-bp period in the *Xoo* PXO83 genome. (**e**) For a large DNA repeat region with a repeat unit size of 787 that overlaps with three different hypothetical genes in the *Xoo* PXO83 genome.

Except for TALEs, ~88% (156 out of 178) of protein repeats were composed of short tandem repeats with a repeat unit size of 10 aa or less (Supplementary Table 3). The other repetitive proteins included three chemotaxis-associated proteins with different periods of 27, 46, and 90 aa, a DNA topoisomerase I, a TolB-like protein known to involve non-WD40 β-propellers, and transporters and a hypothetical protein involving six repeats with a large unit size of 215 aa. Notably, another type III secretion system effector protein of the *Xanthomonas* host infection process was found to have repetitive peptide units, suggesting another function of pathogenic periodic protein structure in hijacking the host plant system (Supplementary Fig. 4).

## Performance in detecting TALEs and C2H2 ZFNs

As C2H2 ZNFs are the most widely used transcription factors in the human genome, we also examined whether SPADE can capture human C2H2 ZNFs. When a C2H2 ZNF encoded on human chromosome 7p22.1 was scanned by SPADE, 20 degenerative repeats of ~28 aa were detected with two cysteine and two histidine residues conserved at specific positions, like typically reported C2H2 ZNFs (Fig. 4). We then assessed the performance of SPADE in detecting TALEs and C2H2 ZNFs. Using the protein domain search software HMMER, we obtained positive reference sets (PRSs) for TALE and human C2H2 ZNF from the prokaryotic genomic dataset and the human proteome, respectively, so each PRS protein contained three or more of the corresponding Pfam motifs (see Methods). We also prepared 10,000 prokaryotic proteins and 10,000 human proteins that did not have any Pfam motif more than once as negative reference sets (NRSs) ProNRS10K and HuNRS10K, respectively (see Methods). Using SPADE, repetitive sequences of any period were detected in 328 out of 331 TALE PRS proteins (99.1%) and 3,079 out of 4,084 human C2H2 ZNF PRS proteins (75.4%), while 192 ProNRS10K proteins (1.9%) and 1,269 HuNRS10K proteins (12.7%) were positive (Fig. 5a and Supplementary Table 4). When the detected positives were filtered by maximum repeat unit size per protein (maxRUSPP) to be within ±5 aa from the expected average repeat unit size (34 aa for TALE and 28 aa for C2H2 ZNF), the recall of TALE PRS stayed the same (99.1%) and the recall of human C2H2 ZNF PRS was 58.9%, while the false positive rate (FPR) of TALE estimated using ProNRS10K and the FPR of C2H2 ZNF estimated using HuNRS10K were greatly decreased to 0.03% and 0.21%, respectively (Fig. 5b and c). This simple size limitation improved positive likelihood ratios (PLRs) of the prediction from 51.6 to 3,303.1 (64.0-fold) for TALE and from 5.9 to 280.7 for human C2H2 ZNF (47.2-fold).

**Figure 4.**
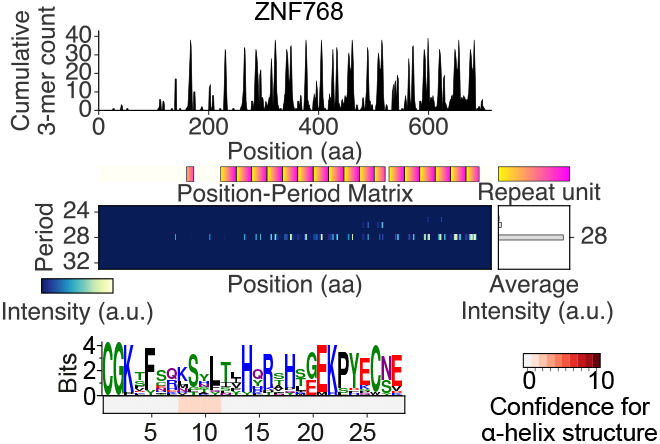
Periodic features of human ZNF768 detected by SPADE.

**Figure 5.**
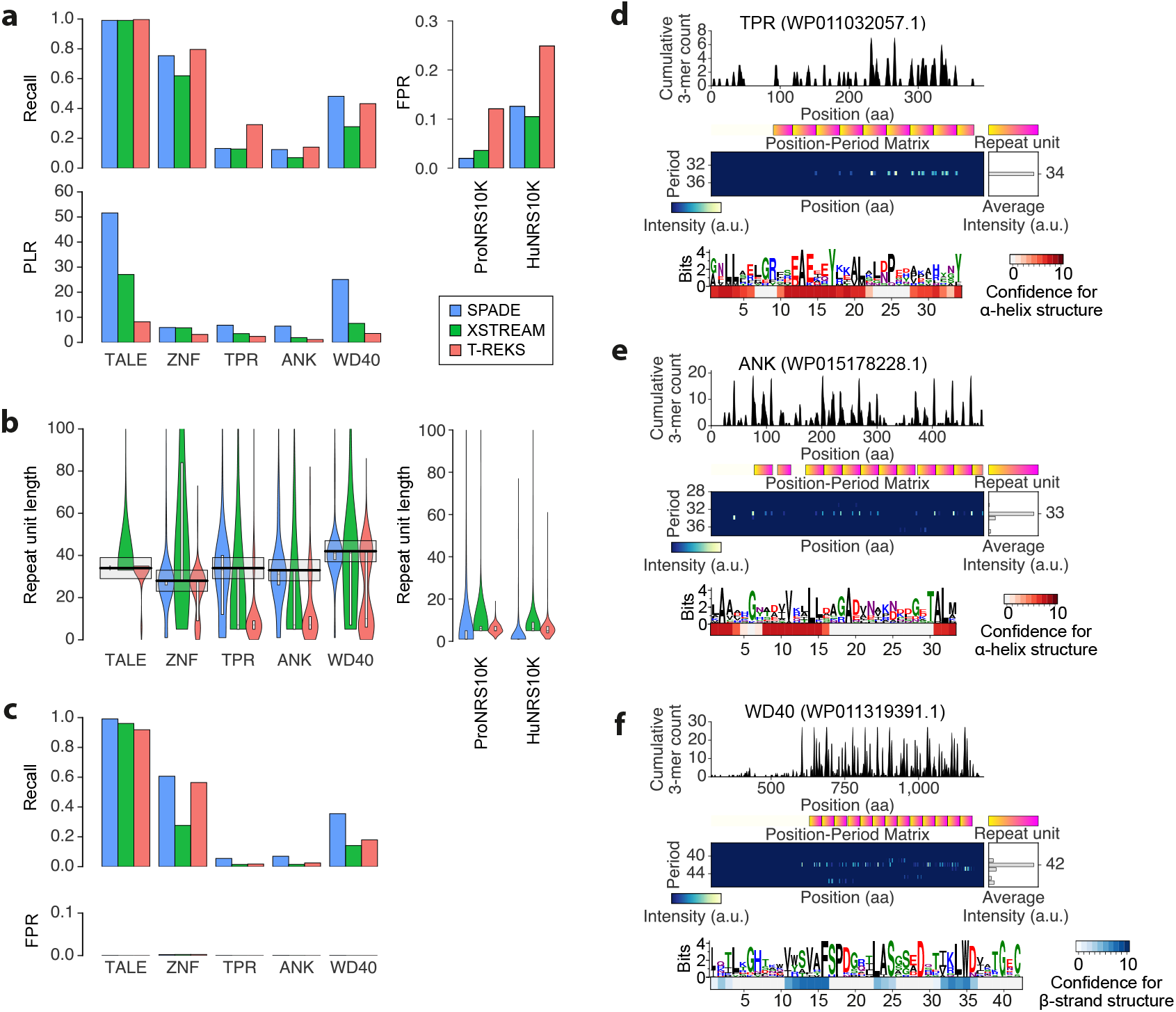
Comparison of the performance of SPADE, XSTREAM, and T-REKS to detect tandem degenerate protein repeats. (**a**) Recalls, false positive rates (FPRs), and positive likelihood ratios (PLRs) of SPADE, XSTREAM, and T-REKS in capturing TALEs, ZNFs, TPRs, ANK repeats, and WD40 repeats. (**b**) Distribution of maximum repeat unit sizes per protein (maxRUSPPs) detected by each software for each protein family. Each bold black bar and each gray box denote the reported typical repeat unit size for the corresponding protein category and the ±5 aa range from the reported repeat unit size, respectively. (**c**) Recalls and FPRs of the different software after filtering maxRUSPPs of the detected positives to be within ±5 aa from the expected repeat unit size. From the maxRUSPP distributions of ProNRS10K (negative control for prokaryotic protein repeat families) and HuNRS10K (negative control for ZNF s), FPRs for different protein repeats of different expected repeat unit sizes were estimated. (**d**–**f**) Example protein repeats detected by SPADE for TPR (**d**), ANK repeat (**e**), and WD40 repeat (**f**). The heat map under each repeat motif sequence logo represents the confidence scores for α-helical structure (red heat map) or β-sheet structure (blue heat map) at each amino acid residue position.

## Comparison with other software capturing tandem protein repeats

SPADE successfully detected the other degenerate tandem protein repeats widely spread in prokaryotes, including TPRs, ANK repeats, and WD40 repeats (Fig. 5d–f). The secondary structure prediction of these degenerated repeats also properly captured their reported structural motifs. In each of the repeat sequence motifs identified for a TPR and an ANK repeat, both of which have been reported to have helix–turn– helix structures, we observed two α-helical loops (Fig. 4d and e). Four β-strands were also captured in a repeat sequence motif of WD40, consistent with its β-propeller structure (Fig. 4f). As XSTREAM^34^ and T-REKS^35^ have been widely used to explore tandem protein repeats in an unsupervised manner in recent studies^43,44^, we next performed a benchmark comparison of SPADE, XSTREAM, and T-REKS in detecting TPRs, ANK repeats, and WD40 repeats, in addition to TALEs and human C2H2 ZNFs. For TPRs, ANK repeats, and WD40 repeats, PRSs were prepared as described above for TALE. ProNRS10K and HuNRS10K were again used as NRSs for detecting repeats in prokaryotic and human protein families, respectively.

T-REKS performed the best in recall for detecting repetitive sequences regardless of repeat unit size, except for WD40, in which SPADE performed the best (Fig. 5a and Supplementary Table 4). However, T-REKS also demonstrated the highest FPRs in both ProNRS10K and HuNRS10K datasets. When the overall prediction performance was estimated by PLR, SPADE performed the best in every repeat type (between 1.02-fold and 3.39-fold compared with the second-best software XSTREAM for all repeat types). We also found that the maxRUSPPs detected by SPADE were distributed with peaks at 34, 33, and 42 aa for TPR, ANK repeats, and WD40 repeats, respectively, all of which were the reported typical unit sizes for these protein repeats (Fig. 5b). This was not the case for all of the repeats detected by XSTREAM and T-REKS. XSTREAM captured wider ranges of repeat unit sizes for every repeat type and T-REKS tended to capture shorter tandem repeats for the subpopulation of positive reference proteins for TPRs, ANK repeats, and WD40 repeats. Filtering the detected positives by maxRUSPP to be within ±5 aa from the expected average repeat sizes, the recall performance of SPADE was the best for all repeat types, whereas the FPRs of the three software packages were all minimized to below 0.005 in all of the repeat types (Fig. 5c and Supplementary Table 4). (Note that the performances could not be compared using PLR as many FPRs for different protein families were zero.) These observations were maintained when the positive reference protein sets were prepared differently (Supplementary Fig. 5).

## Discussion

We demonstrated that SPADE could detect various periodic biomolecular sequences. No software programs have been developed that can universally screen for periodic DNA and protein repeats; the only available software tools are those that screen for reported motifs or certain types of periodic repeats. Nevertheless, the performance of SPADE capturing CRISPRs was on par with the commonly used CRISPR prediction software CRISPRFinder and outperformed XSTREAM and T-REKS in the sensitivity for capturing various tandem protein repeats, regardless of the degree of consensus in the repeat unit motifs. SPADE also captured TALEs and ZNFs in a highly specific and unsupervised manner, indicating its potential to contribute to the discovery of new genome editing modules from large genomic and/or metagenomic resources. This is supported by the fact that we found that a non-TALE type III secretion system protein of *Xanthomonas* host infection machinery had periodic repeats like TALEs and ZFNs (Supplementary Fig. 4). We also captured bacterial homologs of pentatricopeptide repeats (PPRs) that are involved in translational regulation in plants (Supplementary Fig. 6). As the binding code of PPR to RNA has recently been deciphered, it has been suggested as a potential programmable RNA regulating module^45^.

The majority of the periodic repeats detected in the 7,006 prokaryotic genomes still need further investigation. We detected many short tandem DNA and proteins repeats. In particular, tandem heptamer DNA repeats were the most abundant in intergenic regions of a wide range of prokaryotic species (Fig. 1d). However, there has been no clear clue about the function of this globally existing prime number periodicity in genomic DNA. We also found various interspaced repeats that had clear sequence periodicities with no CRISPR annotation or neighboring Cas gene. They included many tRNA operons in various prokaryotes, as reported previously (Supplementary Fig. 7), but the others remain to be explored. Genomic expansion and contraction have been thought to occur at the tandem repeat sequences, leading to phase variation. Even after excluding corresponding protein repeats, the repeat periods of both tandem and interspaced DNA repeats showed particular abundance for these in multiples of three. Furthermore, some genes were encoded in part of a repeat unit of a large tandem repeat region (Supplementary Fig. 3). As seen in the *Xanthomonas* genus (Fig. 3e), these findings suggest the roles of tandem repeats in *de novo* gene birth or gene death. We also found many tandem DNA repeats within (or partially within) protein-coding regions, some of which were indicated to have contributed to functional phase variation of protein-coding patterns (Fig. 3a–d).

SPADE was implemented using Python and can be executed on MacOS X and Linux operating systems with MAFFT and BLAST installed. Default parameters can be used to robustly capture most of the repeats, as demonstrated. It automatically detects the input file and sequence types and outputs results in the GenBank file format, which can be further input into other software programs, with various visualizations as represented in the figures. SPADE precisely captured most of the important biological sequences tested in this study with higher precision than did the other software. Accordingly, here we propose that SPADE is fast and user-friendly software based on a simple algorithm to globally capture periodic biomolecular sequences. Although we mainly focused on measuring the performance of this software predominantly using prokaryotic genomes in this study, further wide-ranging investigations of these periodically repeating sequences together with screening of eukaryotic and metagenomic resources could lead to the discovery of new biological events and genome editing tools.

## Acknowledgments

We thank the members of the Yachie laboratory at The University of Tokyo and Keio University for useful comments and discussions throughout the course of this study. We also sincerely appreciate Hirotada Mori for sharing his laboratory space at the Nara Institute of Science and Technology (NAIST). This study was mainly funded by the New Energy and Industrial Technology Development Organization (NEDO) and partly supported by Japan Society for the Promotion of Science (JSPS), Japan Science and Technology Agency (JST) PRESTO program, Japan Agency for Medical Research and Development (AMED) PRIME program, The Naito Foundation, The Nakajima Foundation, The Takeda Foundation, and SECOM Science and Technology Foundation (to N.Y.). It was also supported by TTCK fellowships (to H.M., D.E.-Y., and S.I.), the Mori Memorial Foundation (to H.M.), the Yamagishi Student Project Support Program (to D.E.-Y.) of Keio University, and a JSPS DC1 Fellowship (to S.I.).

## Author Contributions

H.M. and N.Y. conceived and designed the whole study and wrote the manuscript. H.M. implemented the software. H.M. performed all of the data analyses. D.E.-Y., S.I., and M.T. provided crucial discussions on the software design and data analysis.

## Methods

### Software availability

The SPADE software package and the instruction manual are available at https://github.com/yachielab/SPADE.

### SPADE screening phase 1: Detection of highly repetitive regions

In SPADE, each entry sequence is first scanned by a sliding window of a given size *w* to roughly detect highly repetitive regions (HRRs). Let *W_i_* and *k_i_* be the sliding window at sequence position *i* of the entry sequence and its left-most *k*-mer sequence, respectively. At every sliding window position *i*, the number of *k_i_* within *W_i_* (*n_i_*) is cumulatively counted for every position in the left-most *k_i_* sequence region. In the same sliding window, 1 is also counted for every position of the other *k_i_* sequence regions. Let *c_i_* be the resulting cumulative *k*-mer score at position *i* of the entry sequence. After scanning the query sequence with the sliding window, cumulative *k*-mer peak areas (CKPAs) for which the peak heights are *s* or more are extracted. From the sequence regions for all of the CKPAs (*c* > 0), the broadest possible regions consisting of multiple gaps of size *g* or less were extracted as highly repetitive regions (HRRs). We adopted *w* = 1,000, *k* = 10, *s* = 20, and *g* = 300 for nucleotide sequence and *w* = 300, *k* = 3, *s* = 6, and *g* = 50 for protein sequence as default parameters of the software.

### SPADE screening phase 2: Evaluation of periodicity

Let *h* be the size of a given HRR. For each HRR surrounding region of size *h* ± m, SPADE generates a position-period matrix (PPM) using a similar sliding window of size *h*. Let *Wi* and *k_i_* be the sliding window at sequence position *i* and its left-most *k*-mer sequence, respectively. When multiple *k_i_* sequence regions are detected in *W_i_*, the number of *k_i_* regions (*n_i_*) is cumulatively counted for all of the corresponding row–column cells of the first two k-mer regions, where row represents distance between two identical *k*-mers and column represents sequence position. When *n_i_* > 2, from the second *k_i_* sequence regions, this procedure is iteratively repeated except that the number added to each cell is 1. After scanning by the sliding window, the highest peak period *d* in the column sum distribution of the resulting PPM is identified. All values in a sub-PPM of rows from [*d* × 0.8] to [*d* × 1.2] are then added up and divided by that from 1 to half the column size of the PPM to produce the periodicity score. HRRs with periodicity scores of *p* and more are redefined as periodic repeat regions (PRRs). We set *m* = 1,000 and *p* = 0.5 for nucleotide sequence and *m* = 300 and *p* = 0.3 for protein sequence as default parameters.

### SPADE screening phase 3: Identification of repetitive motifs

From each PRR with sequence period *d*, the *k*-mer sequence that has contributed the most to the sequence periodicity is extracted as *k*seed. When multiple *k*-mer sequences are extracted as the *k*-mers contributing the most, the left-most *k*-mer in the PRR is selected as *k*_seed_. Starting from all of the k_seed_ sequencs found in the PRR, SPADE obtains sequence fragments of size *d*. The extracted sequences are then aligned by multiple sequence alignment using MAFFT version 7.22^46^ to identify their consensus sequence motif. For each sequence position of the alignment result, the information content of appearing letters (*b*, bit) and the frequency of alignment gaps (*f* are calculated using the Python WebLogo 3.6.0 package^47^. After removing positions with *f* of more than *q* from the alignment result, letter consistency *l* of every position is calculated by *b × f*. The positions of the alignment result are then treated as circular since they are for periodic repeats and punctuated by removing the longest continuous nonconsensus region (*l* < *u*) of more than *r* letters. When this punctuation does not happen, the sequence alignment result is linearized as it was before. To map the repeat motif to the PRR sequence, a representative sequence is obtained by taking the most frequent letter in each position of the alignment result. When the representative sequence is shorter than *k*-mer, the identical sequence regions are scanned in the PRR and annotated as repeat units. Otherwise, the representative sequence is mapped using BLAST+ version 2.6^48^ with the blastn-short (for nucleotide) or blastp-short (for protein) option and alignment length threshold of 50% to the query length or E-value of 0.01 or less. The hit regions in the PRR are then used to construct a sequence logo profile using Python WebLogo 3.6.0 package^47^. From the sequence logo profile, a repeat motif sequence is generated by the most frequent letters, where a highest letter frequency of less than 60% is masked with ‘*’. *q* = 0.5, *u* = 0.8, and *r* = 5 were set as default parameters of the software.

### Protein secondary structure prediction

For each visualized protein repeat motif sequence, the confidence score for α-helix structure or β-sheet structure was calculated using PSIPRED version 3.3^49^. For each PRR detected by SPADE, PSIPRED was initially used to predict all possible secondary structure motifs with the confidence score at each amino acid residue position. We then calculated the average confidence score for each motif at every position in the repeat sequence unit.

### Genomic resources

The GenBank files for the 7,006 complete prokaryotic genomes (downloaded on March 31^st^, 2017) and the human reference genome version GRCh38.p10 were downloaded from the NCBI RefSeq genomes FTP server (ftp://ftp.ncbi.nlm.nih.gov/genomes/refseq/).

### Evaluation of performance for detecting CRISPRs

From the entire periodic DNA repeats detected by SPADE, we extracted CRISPR candidates with interspace sizes of 25–60 bp and repeating periods of 58–81 bp. The interspace size parameters and the minimum threshold for the repeating period (interspace size plus repetitive sequence size) were set with reference to the CRISPRFinder screens for CRISPR candidates with interspace size being 25–60 bp and repetitive sequence size being 23–55 bp, but our maximum threshold for the repeating period was defined empirically based on the reported RefSeq CRISPRs. Region overlap agreement (ROA) between two given regions was calculated by dividing the size of the overlapping region by the combined size of the two regions. Recall and precision of the recapturing RefSeq CRISPRs were evaluated for each ROA threshold.

### Evaluation of performance for detecting tandem protein repeats

From the 7,006 prokaryotic genome resources, we screened the positive reference set (PRS) proteins for TALE, TPR, ANK repeat, and WD40 repeat families using HMMER version 3.1 with the Pfam domain signatures of PF03377, PF00515, PF00023, and PF00400, respectively. The PRS proteins for the C2H2 ZNF family were screened from the human reference genome version GRCh38.p10 using the Pfam domain signature of PF00096. Every PRS protein was required to contain three or more of the corresponding Pfam domain copies mapped with an E-value of less than 1.0e–10, and we obtained 331, 26,289, 4,428, 2,672, and 4,084 PRS proteins for TALE, TPR, ANK repeats, WD40 repeats, and C2H2 ZNF, respectively (Supplementary Table 4). A total of 100,000 randomly picked prokaryotic proteins and the entire human proteome were screened for Pfam-A domain families version 31.0. Among those that do not have more than one copy of any Pfam domain with an E-value of less than 1.0e–10, we randomly selected 10,000 prokaryotic proteins and 10,000 human proteins as negative reference sets ProNRS10K and HuNRS10K. The performance of the software programs SPADE, XSTREAM, and T-REKS was estimated using the recall of PRS proteins and the false positive rate (FPR) in ProNRS10K (for prokaryotic protein repeats) or HuNRS10K (for human protein repeats). The positive likelihood ratio (PLR) was calculated by dividing recall by FPR. Each software was used with its default parameters. Similar analysis was also performed by restricting the detected repeat unit sizes to within the range of expected sizes for different repeat families (34±5 aa, 34±5 aa, 33±5 aa, 42±5 aa, and 28±5 aa for TALE, TRP, ANK repeats, WD40 repeats, and C2H2 ZNF, respectively). Note that, owing to the size filtering, FPRs varied for different repeat families, even when the same negative reference set was used. These measurements were also repeated with PRS proteins prepared using different criteria (Supplementary Fig. 5).

